# Ca^2+^ Regulation of Myosin II and Myosin VI during Rupture of the *Shigella*-Containing Vacuole

**DOI:** 10.1101/2024.05.22.595272

**Authors:** Mariette Bonnet, Chun Hui Sun, Jost Enninga, Anne Houdusse, Guy Tran Van Nhieu

## Abstract

*Shigella*, the causative agent of bacillary dysentery, invades epithelial cells to colonize the intestinal mucosa. Following invasion, *Shigella* is enclosed in a vacuole that needs to rupture for bacterial intra-cytosolic replication. We show here that rupture of the *Shigella* vacuole requires Ca^2+^ influx leading to long lasting local Ca^2+^ increases that regulate actin dynamics affecting the *Shigella* vacuole integrity. These Ca^2+^ increases promote vacuolar rupture by activating myosin II associated with actin filaments in membrane ruffles distant from the vacuole, while tethering myosin VI at the actin coat-surrounded vacuole. Ca^2+^ depletion and myosin II inhibition impair formation of the actin coat and vacuole rupture. Inhibition of myosin VI also delays rupture of vacuoles but lead to their tumbling. These findings highlight a role for Ca^2+^ in coordinating actin–based forces and constraints during early rupture steps of bacterial vacuole, that pull on vacuolar membranes and tether them to the actin cortex via myosin II and VI, respectively, a process relevant to intracellular pathogen and endomembrane trafficking.

## Introduction

*Shigella* is a highly infectious pathogen responsible for bacillary dysentery in humans, and a major cause of endemic diarrheal disease associated with high mortality rates in developing countries (Baker and The, 2018; Kotloff, 2019). Upon ingestion, this bacterial enteropathogen invades the colonic intestinal mucosa where it induces a strong inflammatory response responsible for the destruction of the epithelium. *Shigella* invasion of epithelial cells is associated with actin reorganization of the cellular cortex leading to membrane ruffling at the bacterial invasion site and the formation of a transient bacterial adhesion structure at the bacterial-cell contact. Once internalized by the cell, *Shigella* rapidly escapes from the vacuole to replicate freely in the host cell cytosol (Kuehl et al., 2015; Schnupf and Sansonetti, 2019). During this intracellular replication phase, *Shigella* uses actin-based motility to propel itself into the cytosol and to form protrusions invading adjacent cells, enabling its dissemination in the intestinal epithelium (Kuehl et al., 2015; Agaisse, 2019). *Shigella* is unable to replicate in the vacuole and its escape from it determines bacterial dissemination in the epithelial tissue.

The virulence of *Shigella* requires a so called type three secretion system (T3SS) or “injectisome”, allowing the translocation of at least 30 bacterial effectors into the cytosol of host cells upon cell contact (Schnupf and Sansonetti, 2019; Muthuramalingam et al., 2021). Injected type III effectors divert cellular processes to promote tissue colonization (Muthuramalingam et al., 2021) and can be divided in two main groups based on the regulation of their expression that also determines their role in the infectious process (Schnupf and Sansonetti, 2019; Sanchez-Garrido et al., 2022; Killackey et al., 2016). Constitutively expressed type III effectors play an early role during *Shigella* interaction with epithelial cells and include the bacterial invasion effectors. Type III effectors that are up-regulated upon cell contact have been involved in the diversion of cellular immunity, cell death or inflammatory processes. Rupture of the *Shigella* containing vacuole (SCV) requires the T3SS apparatus as well as the joint action of several of its effectors. The IpaB and IpaC components of the type III translocon, were initially proposed to trigger vacuolar rupture (Bianchi and Van den Bogaart, 2020). However, the T3SS translocon is also required for the translocation of injected type III effectors, questioning whether its role in vacuolar rupture is direct or indirect. More recent works indicated the involvement a number of invasion type III effectors demonstrating that vacuolar rupture is a multistep process and that the invasion and vacuolar rupture processes are associated events. *Shigella* strains deficient for the type III effectors IpgB1 and IpgB2, acting as GEFs for the small GTPases Rac and Rho, respectively, show a delay in vacuolar rupture suggesting a role for actin dynamics (Mellouk and Enninga, 2016). The type III invasion effector IpgD, a phosphatidylinositol (PI) 4, 5 biphosphate (PtdIns(4,5)P_2_) - 4 phosphatase, also recently characterized as a phosphotransferase / isomerase generating (PtdIns(3,4,5)P_3_) and (PtdIns(3,4)P_2_) from (PtdIns(4,5)P_2_), was also shown to contribute to SCV rupture (Mellouk et al., 2014; Walpole et al., 2022; Tran Van Nhieu et al., 2022). IcsB is another T3SS effector acting on host GTPases through N-fatty acylation that affects vacuolar rupture by altering the actin coat forming around the SCV before rupture (Kuhn et al., 2020). Furthermore, the effector IpaH7.8 is not involved in early vacuolar damages but has been shown to promote the full unwrapping of damaged SCVs along microtubules (Chang et al., 2024).

Live imaging microscopy analysis indicates that *Shigella* vacuolar escape occurs within minutes following bacterial internalization in the continuation of the invasion process (Ehsani et al., 2012). However, while *Shigella* invasion has been largely studied and general features of its underlying mechanism are reasonably understood, much less is known about the precise nature of the forming SCV and the mechanism leading to its rupture. *Shigella*-induced membrane ruffles during invasion are associated with the formation of macropinosomes which are distinct from and do not fuse with the SCV (Weiner et al., 2016). Macropinosomes appear to participate in vacuolar rupture because their tight apposition with the SCV correlates with its loss of integrity (Weiner et al., 2016). These macropinosomes are associated with endosomal components, such as the small GTPase Rab5 and Rab11 and also exocyst components like Exo70, and their depletion delays vacuolar rupture (Mellouk et al., 2014; Weiner et al., 2016; Chang et al., 2020). Furthermore, the dynein complex has been found on the macropinosomes, and broken SCVs are pulled away from the bacterium along microtubules (Chang et al., 2024). Together, these studies have revealed that SCV rupture takes place through subsequent steps including its initial damage, followed by SCV rupture and the later step of complete SCV membrane remnant unwrapping away from the bacteria.

Here, we analyzed the role of Ca^2+^ fluxes during the early steps of *Shigella*-induced vacuolar rupture. We show that long lasting local Ca^2+^ increases associated with influx play a role in this process. We further demonstrate two distinct effects of these Ca^2+^ increases, one through the activation of distally located myosin II and another one on myosin VI that is present at the SCV. Upon Ca^2+^ increase, Myosin VI co-opting immobilizes the SCV within the cortex allowing the pulling on its membranes by distal Myosin II and rupture of the *Shigella*-containing vacuole.

## Results

### Long lasting local Ca^2+^ increases associate with *Shigella* vacuolar rupture

*Shigella* invasion of epithelial cells triggers atypical long lasting Ca^2+^ increases termed Responses Associated with Trespassing Pathogens (RATPs), that are dependent on inositol 1,4,5, -triphosphate (IP3) (Tran Van Nhieu et al., 2013). Imaging of local Ca^2+^ increases is generally not compatible with combined dual imaging due to the required speed of image acquisition in the order of tens of milliseconds. The long duration of RATPs, however, enabled us to visualize local Ca^2+^ increases associated with *Shigella* invasion, while imaging vacuolar rupture using the Galectin-3 lectin (Gal3) (Methods; Paz et al., 2010). This technique enables to detect within seconds the overall integrity of the SCV. The onset of actin foci formation induced during *Shigella* invasion was the reference starting time point set up to precisely measure the time associated with rupture (Fig. 1A).

**Figure 1.**
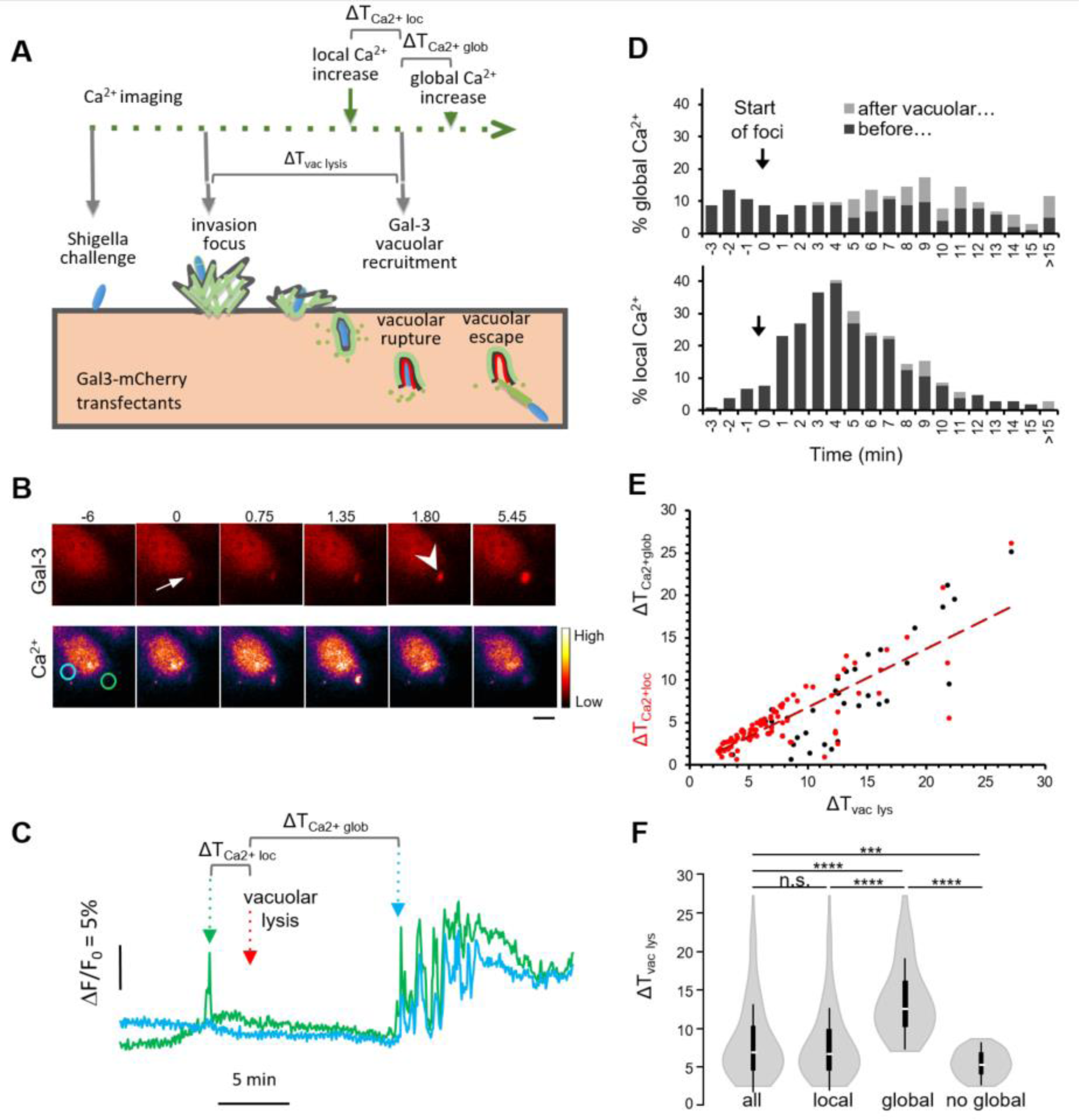
Long lasting local Ca^2+^ increases associate with *Shigella* vacuolar rupture. Gal3-mOrange transfected cells were loaded with Fluo4-AM and challenged with *Shigella*. Ca^2+^ and vacuolar lysis imaging were performed by following the Fluo4 and mOrange fluorescence every 3 seconds, acquiring z-stacks to cover the cell volume for each time point (Methods). The dynamics of invasion foci formation were monitored by following Gal3-mOrange recruitment (Methods). **A**, Scheme of dual Ca^2+^ and vacuolar rupture imaging. The time points corresponding to the start of actin formation and vacuolar rupture were determined by quantifying the Gal3-mOrange average fluorescence intensity at invasion sites (Fig. EV1; Methods). The kinetics of vacuolar rupture were analyzed by determining the time interval between the start of actin foci formation and vacuolar lysis and (ΔT_vac lys_), the shortest time interval between vacuolar rupture and a preceding or following local (ΔT _Ca_^2+^ _loc_) or global Ca^2+^ (ΔT _Ca_^2+^ _glob_) increase. **B**, Representative time-series of fluorescence micrographs. Numbers: time relative to the start of actin foci formation in min. Scale bar = 5 μm. Gal3: maximal z-projection of Gal3-mOrange planes. Arrow: start of invasion foci. Arrowhead: vacuolar rupture. Ca^2+^: Fluo-4 displayed in fire pseudo-color with the corresponding fluorescence intensity scale. **C**, Traces corresponding to variations in intracellular Ca^2+^ concentrations in the cell region as depicted in panel B with matching color. **D**, percent of cells showing global (top) or local (bottom) Ca^2+^ increases preceding (grey) or following (black) vacuolar rupture at the indicated time point. Arrow: the reference time point was set as the start of the first invasion foci for each cell. **E**, time interval between local (black, ΔT _Ca_^2+^ _loc_) or global (red, ΔT _Ca_^2+^ _glob_) Ca^2+^ increases plotted as a function of the time interval of vacuolar rupture (ΔT_vac lys_). Solid line: linear fit for ΔT _Ca_^2+^ _loc_ over ΔT_vac lys_. Pearson coefficient: 0.81. **F**, violin plot showing ΔT_vac lys_ in cell subgroups corresponding to all cells analyzed (all), or cells showing only local, global, or no global Ca^2+^ increase. n = 107 cells, N = 7. Mann-Whitney test. ns: not significant; ***: p <0.005; ****: p < 0.001.

In a first set of experiments, cells were transfected with GFP-actin and Gal3-mOrange and their recruitment kinetics at *Shigella*-induced entry sites were quantified (Fig. EV1A, dotted cyan and orange ROIs, respectively). As shown in Fig. EV1B, the onset of actin polymerization could be readily determined as the inflexion in the mean intensity trace of GFP-actin (Fig. EV1B, cyan arrow). We observed that the onset of actin foci formation also matched an inflexion in the mean intensity of Gal3-mOrange recruitment at *Shigella*-induced foci (Fig. EV1B, orange arrow). This increase in Gal3-mOrange intensity likely corresponded to the accumulation of membrane ruffles formed at bacterial entry sites and could be clearly distinguished from the sharp increase in Gal3-mOrange intensity associated with vacuolar rupture when imaging Gal3-mOrange recruitment at the SCV (Fig. EV1B, dotted red ROI and red arrow). We concluded from these experiments that imaging of Gal3-mOrange could be simultaneously used to follow *Shigella*-induced foci formation during bacterial invasion and vacuolar rupture, and could be advantageously combined with Ca^2+^ imaging to limit photo-damage associated with multiple wavelength acquisition.

As illustrated in Figs. 1B, C, a local Ca^2+^ increase could be readily detected in a region corresponding to the site of bacterial-induced foci (Fig. 1B, green ROI and white arrow, respectively) preceding vacuolar rupture by 12-15 seconds (Fig. 1B, white arrowhead; Movie EV1). Global Ca^2+^ increases were also observed within a few minutes following vacuolar rupture (Fig. 1C). In Figs. 1D, E, we reported the time intervals measured for local and global Ca^2+^ increases, differentiating between those elicited before and after vacuolar rupture. As shown in Fig. 1D, global Ca^2+^ increases were detected during the first 15 minutes following the formation of *Shigella*-induced invasion foci in the absence of obvious correlation with vacuolar rupture. In contrast, local Ca^2+^ responses predominantly occurred during the first five minutes following foci formation, with a remarkable correlation with vacuolar rupture (Fig. 1E). This correlation stood out when plotting the time interval of local Ca^2+^ responses relative to vacuolar rupture, as a function of the time interval of vacuolar rupture (Fig. 1F, ΔT_Ca2+loc_, red dots), whereas no correlation was observed for global Ca^2+^ responses (Fig. 1F, ΔT _Ca_^2+^_glo_, black dots). Interestingly, when specifically analyzing sub-groups of Ca^2+^-responding cells, we found an inverse relationship between cells showing global responses and the time interval in vacuolar rupture: cells eliciting global Ca^2+^ increases showed longer ΔT_vac lys_ with an average duration of 14.2 min compared to cells showing only local Ca^2+^ increases with an average duration 5.5 min (Fig.1F compare global versus local). Further, cells showing no global Ca^2+^ increase showed even shorter time of vacuolar rupture compared to the other sub-groups (Fig. 1F, no global), suggesting that global Ca^2+^ increase antagonized vacuolar rupture.

These results show a temporal association between long lasting local Ca^2+^ increase and rupture of the *Shigella*-containing vacuole, and suggest that as opposed to global Ca^2+^ increases, the localized aspect of Ca^2+^ increases at the bacterial entry site is a key parameter in favoring vacuolar rupture.

### Ca**^2+^** influx at the bacterial entry site accounts for long lasting local Ca**^2+^** increases and promotes SCV rupture

Local responses associated with vacuolar rupture showed even longer Ca^2+^ increases duration than those observed for RATPs, often exceeding 10 seconds as illustrated in Fig. 1C, suggesting an additional contribution of Ca^2+^ influx. To test this, cells were incubated with 1.8 mM EGTA to chelate extracellular Ca^2+^ prior to bacterial challenge and Ca^2+^ imaging (Methods). As shown in Figs. 2A-C, Ca^2+^ chelation did not affect the percent of cells showing global Ca^2+^ responses induced by *Shigella* (Figs. 2A-C, Global). In contrast, Ca^2+^ depletion drastically reduced the percent of cells showing long lasting Ca^2+^ increases associated with vacuolar rupture (Figs. 2A-C, Local, compare CTRL and No Ca^2+^). Ca^2+^ removal also led to a 2.8–fold increase in the ΔT_vac lys_, indicating a role for Ca^2+^ influx in vacuolar rupture (Fig. 2D).

**Figure 2.**
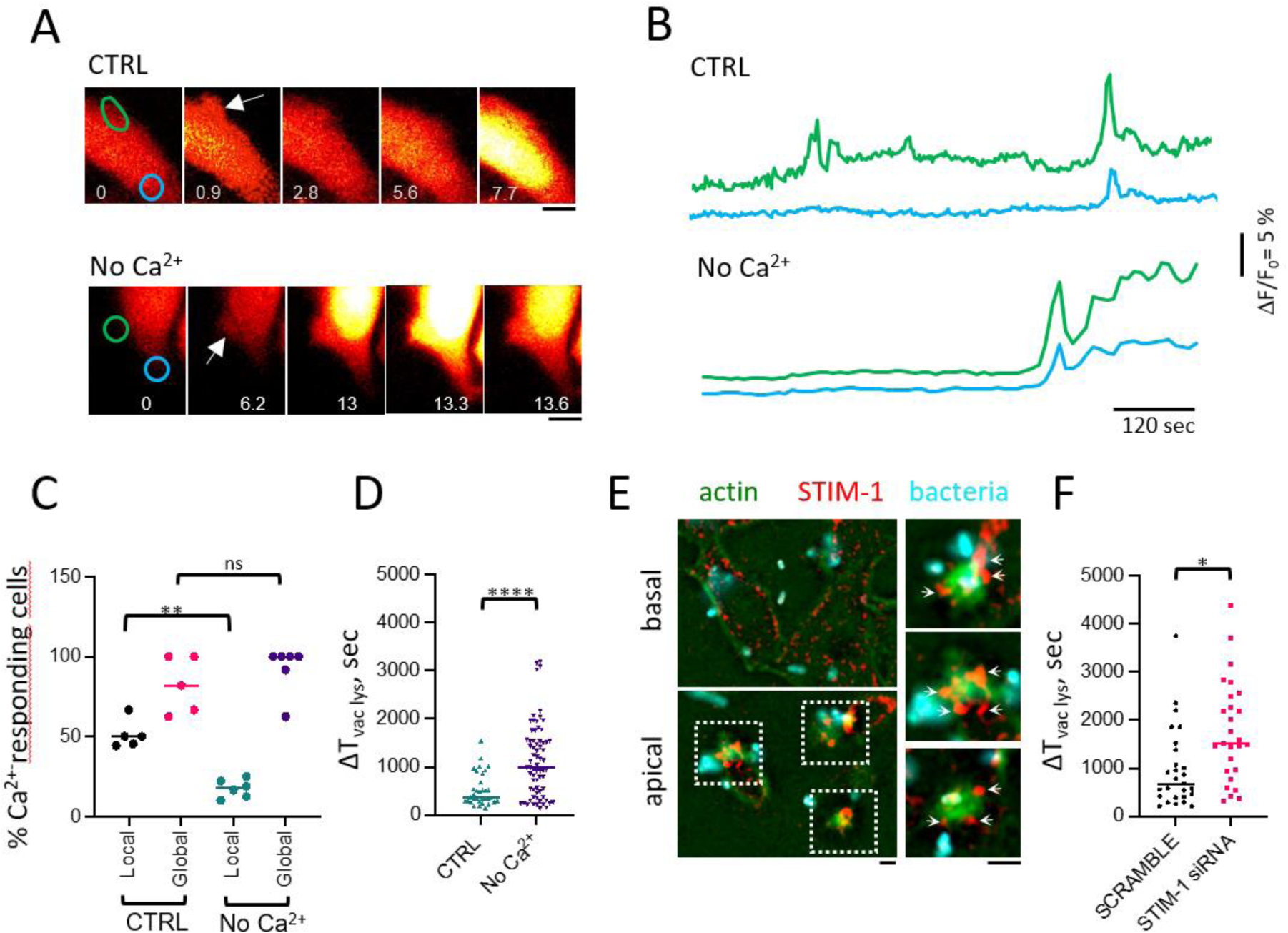
Ca^2+^ influx is required for rupture of the *Shigella*-containing vacuole. Cells were loaded with Fluo4-AM, challenged with *Shigella* in the presence (CTRL) or absence of extracellular Ca^2+^(No Ca^2+^) and subjected to Ca^2+^ imaging. **A**, Representative time-series of micrographs of Fluo-4 fluorescence displayed in fire pseudo-color with the corresponding fluorescence intensity scale. Scale bar = 5 μm. Numbers: time relative to the start of actin foci formation in min. Scale bar = 5 μm. Arrow: invasion foci. **B**, Traces corresponding to variations in intracellular Ca^2+^ in the cell region is depicted in panel B with matching color. **C**, percent of cells showing global and local Ca^2+^ responses in the presence or absence of external Ca^2+^. CTRL: 61 cells, N = 6. No Ca^2+^: 85 cells, N = 5. **D**, Cells were transfected with scramble control (SCRAMBLE) or anti-STIM-1 siRNA (STIM-1 siRNA). Alternatively, cells were treated in the presence or absence of extracellular Ca^2+^. Samples were challenged with *Shigella* and subjected to Gal 3-mOrange based vacuolar rupture imaging (Methods). The ΔT_vac lys_ was determined for the indicated samples. SCRAMBLE: 30 cells, 14 vacuolar rupture events, N = 4. siRNA STIM-1: 58 cells, 27 vacuolar rupture events, N = 5. CTRL: 36 SCVs, N = 3. No Ca^2+^: 69 SCVs, N = 3. Mann-Whitney test. ns: not significant; *: p < 0.05; **: p < 0.01. ****: p < 0.001. **E**, Cells were challenged with wild-type *Shigella* (WT). Representative immunofluorescence staining micrographs of basal and apical planes. Red: STIM-1; green: F-actin; blue: bacterial LPS. Scale bar = 2 μm. Quantitative analysis indicated that STIM-1 showed a 1.5-fold increase in average intensity in invasion foci relative to the rest of the cell. 119 foci, N = 3. Mann-Whitney test. ****: p < 0.001.

*Shigella* recruits ER compartments at invasion foci (Tran Van Nhieu et al., 2013; Sun et al., 2017). This suggests that *Shigella*-induced Ca^2+^ influx is possibly also regulated by ER Ca^2+^ stores involving the ER Ca^2+^ luminal sensor STIM-1 (Stromal Interaction Molecule -1) in a Ca^2+^ release dependent manner. We therefore tested the role of STIM-1 in rupturing the SCV. As shown in Fig. 2E and as previously reported, STIM-1 showed a predominant cytoplasmic punctate staining in control cells (Gudlur et al., 2020; Fig. EV2A). In contrast, STIM-1 showed large clusters at *Shigella* invasion sites forming at the apical surface consistent with its translocation at the plasma membrane and interaction with Ca^2+^ release-activated Ca^2+^ (CRAC) channels (Fig. 2E, arrows and Fig. EV2A, arrowheads). Consistently, we also found the ORAI-1 channel to be recruited at *Shigella* invasion sites (Fig. EV2B, arrowheads).

We next tested the functional implication of STIM-1 in vacuolar rupture by inhibiting its expression by siRNA treatment (Methods). As shown in Fig. EV2E-F, anti-STIM-1 or anti-ORAI-1 siRNAs did not affect the frequency of actin foci formation induced by *Shigella*, relative to control scramble siRNAs. ORAI-1 depletion also did not affect SCV rupture, suggesting that the involvement of other CRAC channels (Fig. EV2G). In contrast, STIM-1 depletion led to a two-fold increase in ΔT_vac lys_ indicating its involvement in rupture of the SCV (Fig. 2F, compare SCR and STIM-1 siRNA).

Together, these results indicate that the long lasting local Ca^2+^ increases associated with formation of the SCV depend on Ca^2+^ influx and contribute to vacuolar rupture.

### Ca^2+^ regulates actin dynamics at *Shigella* invasion sites

Vacuolar rupture predominantly occurs in the continuation of the invasion process, suggesting that actin-based forces exerted during bacterial internalization on the forming vacuole could be involved in the breakage of this latter, possibly in link with the local Ca^2+^ influxes that we observed. Therefore, we performed live imaging measuring the impact of local Ca^2+^ influx on *Shigella*-induced actin dynamics and its role in the steps leading to SCV rupture.

Cells transfected with GFP-actin and Gal3-mOrange were challenged with *Shigella* in the presence or absence of extracellular Ca^2+^ and analyzed by spinning disk confocal microscopy (Methods). As shown in Fig. 3A and as previously described, within a few minutes, *Shigella* induced actin polymerization at the vicinity of the bacterial contact with host cells forming characteristic actin foci in control cells. Concomitantly (Fig. 3A, CTRL-1), or a few tens of seconds later (Fig. 3A, CTR-L2), an actin rich structure, termed “pseudo-focal adhesion” or “actin cup”, formed in tight apposition with the invading bacteria anchoring it into the actin foci (Valencia-Gallardo et al., 2015; Fig. 3A, arrows). As bacterial invasion progressed, this actin coat structure thickened and extended to cover the entire SCV (Fig. 3A; Movie EV2), forming a thick actin coat that has also been named “actin cocoon” (Kühn et al., 2020). Vacuolar rupture generally occurred within 10 minutes following formation of the actin cup or cocoon, with a labeling of the SCV tightly contacting the bacterial body (Fig. 3A, red arrows). At the onset of vacuolar rupture, Gal3 labeling of the SCV often correlated with thinning or partial disappearance of the actin cocoon (Fig. 3A, red arrows).

**Figure 3.**
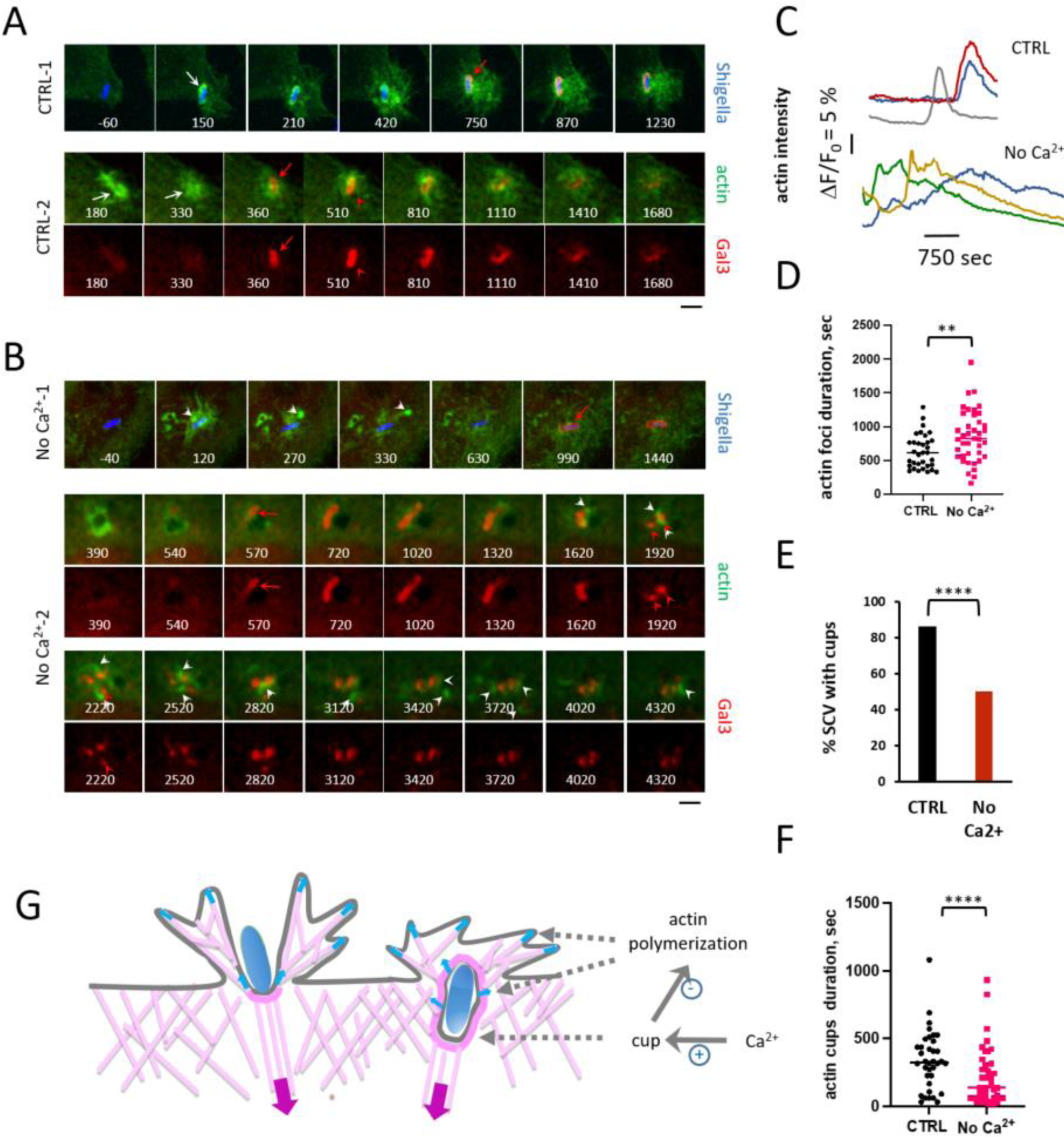
Ca^2+^ regulates actin dynamics at *Shigella* invasion sites. Gal3-mOrange and GFP-actin double-transfected cells were challenged with cerulean-expressing *Shigella* in the presence (CTRL) or absence (No Ca^2+^) of external Ca^2+^. Samples were imaged for GFP-actin and Gal3-mOrange fluorescence by spinning disk confocal microscopy. **A**, **B**, Representative time-series of micrographs of maximal z-projection displayed in fire pseudo-color with the corresponding fluorescence intensity scale. Scale bar = 5 μm. Numbers: time relative to the start of actin foci formation in min. Blue: bacteria; red: Gal3-mOrange; Green: GFP-actin. Scale bar = 5 μm. **A**, **B**, red arrows point at vacuolar rupture. **A**, white arrows: actin cups. **B**, arrowhead: actin-comet structures forming at a distance from the SCV. **C**, representative traces of the average GFP-actin average fluorescence intensity at bacterial invasion site over time. Each color represents an individual invasion focus. **D**, actin foci duration. **E**, percent of SCV with actin cups. CTRL: 58 SCVs, N = 8. No Ca^2+^: 66 SCVs, N = 5. Chi-square. ****: p = 0.000014. **F**, actin cups duration. CTRL: 36 SCVs, N = 8. No Ca^2+^: 32 SCVs, N = 5. Mann-Whitney test. ****: p < 0.001. **G**, scheme of actin dynamics regulation by Ca^2+^ influx at *Shigella* invasion site. Ca^2+^ up-regulates the formation of the actin cup (dark pink) surrounding the SCV and negatively regulates actin polymerization at the plasma and SCV membranes (grey arrows). Light pink: actin filaments. Blue arrows: actin-polymerization based forces exerted at the plasma and SCV membranes. Purple arrow: retrograde flow.

In the absence of external Ca^2+^, while the general features of actin foci, cup and cocoon formation were still observed, a detailed and quantitative analysis indicated major differences associated with a delay in vacuolar rupture. In control conditions, actin foci showed an average duration ± SEM of 10.5 ± 0.7 min (Figs. 3C, D, CTRL). In the absence of Ca^2+^, actin polymerization at invasion sites lasted for a significantly longer periods, with an average duration ± SEM of 14.2 ± 1.0 min (Figs. 3C, D, No Ca^2+^). In control conditions, *Shigella*-induced actin polymerization occurred in membrane ruffles at invasion foci that resorbed after the formation of the cups or cocoons (Fig. 3A, CTRL, Movie EV2). In contrast, in the absence of external Ca^2+^, actin polymerization was often detected for longer periods being visible after vacuolar rupture for extended time periods (Figs. 3B, C, No Ca^2+^; Movie EV3).

Remarkably, extracellular Ca^2+^ also differentially regulated the actin structures (cups and actin cocoon) associated with the SCV. As opposed to control conditions, the majority of invasion events were not associated with actin cup formation without external Ca^2+^, with only 14.7 % compared to 89.1 % of actin cup formation the absence and presence of external Ca^2+^, respectively (Fig. 3E). The remaining actin cups forming in the absence of external Ca^2+^ were more transient, showing a reduced average duration ± SEM of 3.6 ± 0.6 min, compared to control conditions showing an average duration ± SEM of 5.6 ± 0.6 min (Fig. 3F). The transient nature of actin cups and cocoons in the absence of external Ca^2+^, was also associated with specific features of SCV disintegration. While in control conditions, broken SCVs generally remained in tight apposition with invading bacteria for several minutes following rupture, in the absence of Ca^2+^, SCVs often disintegrated rapidly, with remnants propelled by small actin comets observed at the vicinity of invading bacteria (Fig. 3B, white arrowheads; Movie EV3). These events happened earlier than the full unwrapping of SCVs along microtubules (Chang et al, 2024).

Together, these results suggest that Ca^2+^ influx promotes faster kinetics of actin polymerization at *Shigella* invasion sites, while stabilizing the SCV actin cocoon (Fig. 3G). These results are reminiscent of antagonistic processes linking actin polymerization associated with lamellae extension and actomyosin contraction during cell adhesion (Giannone et al., 2004; Lawson and Burrridge, 2014; Shaks et al., 2019) and suggest a role for myosin II activation associated with Ca^2+^ influx in rupture of the SCV.

### Ca**^2+^**-activated Myosin II promotes the early steps of SCV rupture

Earlier studies indicated that myosin II was recruited at *Shigella* invasion sites but was not required for the actual bacterial internalization process (Clerc and Sansonetti, 1987; Rathman et al., 2000). Myosin Light Chain Kinase (MLCK) was shown to play a role in dissemination of *Shigella* from cell to cell, suggesting that myosin II contributed to the engulfment of bacteria-containing protrusions emanating from the donor cell by recipient cell (Rathman et al., 2000). We tested whether myosin II could play a role during the *Shigella* invasion process after entry and before its spread to neighboring cells namely by promoting its release from the SCV upon local Ca^2+^ increase.

To detect the role of Ca^2+^ in myosin activation during *Shigella* invasion, we first performed anti-phospho myosin light chain II immunofluorescence microscopy (Methods) at different time point following bacterial challenge. As shown in Figs. 4A, B, phospho-myosin accumulation was readily detected at *Shigella*-invasion sites 5 min after bacterial challenge (Figs. 4A, B, CTRL). This accumulation was not observed in the absence of external Ca^2+^ consistent with a role for Ca^2+^ influx (Figs. 4A, B, No Ca^2+^). Phospho-myosin II recruitment at *Shigella* invasion sites was still detected after 15 min following bacterial challenge, but with a non-statistically significant difference relative to conditions in the absence of external Ca^2+^ suggesting that myosin II activation occurred transiently (Figs. 3B and S3A). A detailed 3D-analysis of *Shigella* invasion sites indicated that phospho-myosin accumulated in actin rich projections elevating from the bacteria-cell contact site with the cell, but not at the levels of the SCV (Fig. 4C).

**Figure 4.**
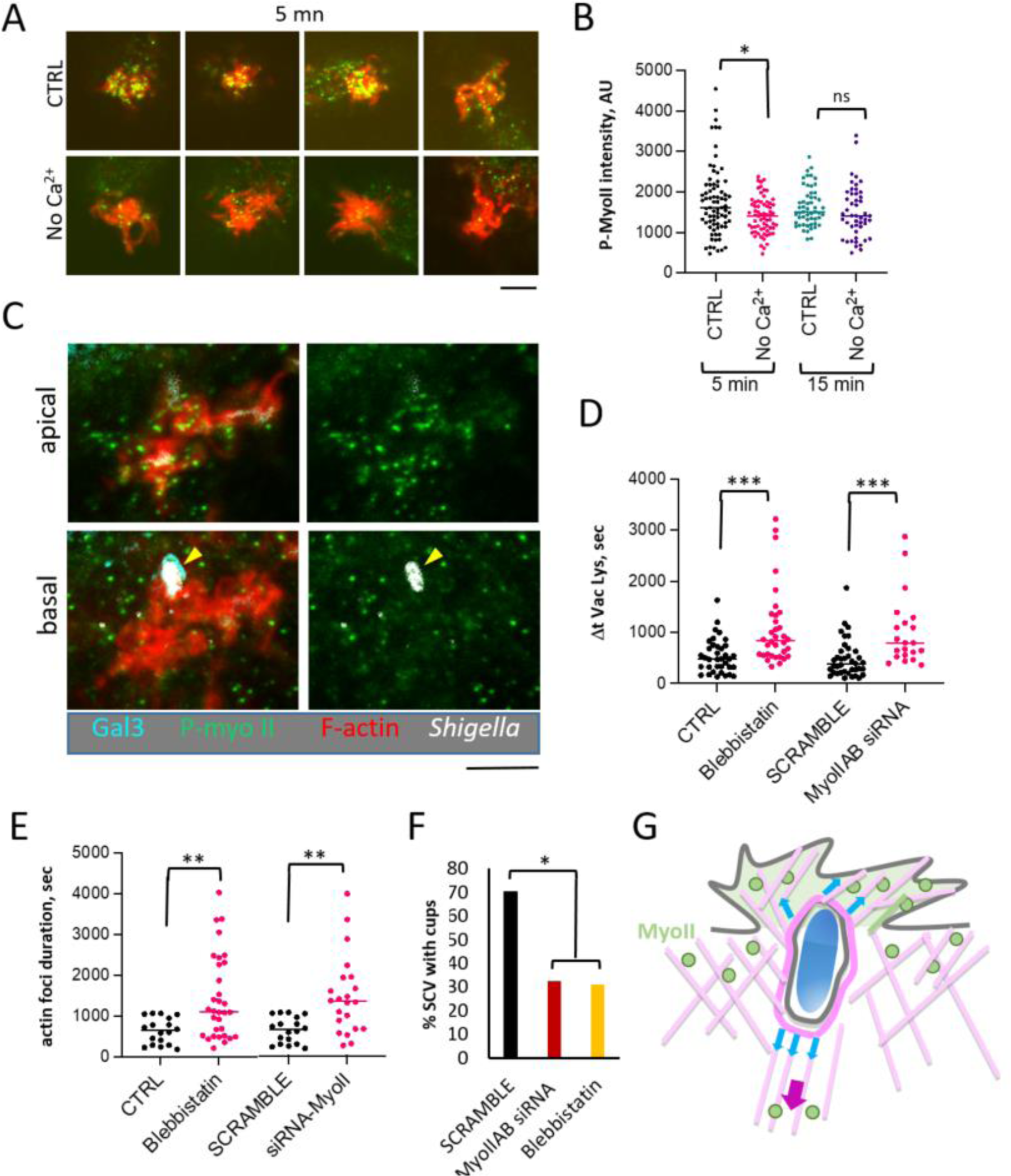
Myosin II is activated by Ca^2+^ and contributes to rupture of the SCV. Cells were challenged with *Shigella*. Samples were fixed and processed for immunofluorescence staining. **A**, **C**, representative fluorescence micrographs. Red: actin; Green: phospho-myosin. Scale bar = 5 μm. **A**, maximal z-projection of confocal planes. Cells infected in the presence (CTRL) or absence (No Ca^2+^) of extracellular Ca^2+^. **C**, cells transfected with Gal3-mOrange prior to bacterial challenge. White: bacterial LPS; cyan: Gal3 labeling of the SCV. Apical and basal confocal plane of *Shigella* invasion site. The yellow arrowhead points at the lack of phospho-myosin staining at the SCV. **B**, quantification of phopho-myosin average fluorescence intensity at *Shigella* invasion sites at the indicated time point. CTRL: 83 foci, N = 6. No Ca^2+^: 73 foci, N = 6. **D**, **E**, control cells (CTRL); cells treated with 50 μM blebbistatin (Blebbistatin), scramble siRNA control (SCRAMBLE); anti-Myosin IIA, B siRNA (MyoIIAB siRNA). Mann-Whitney test. ns: not significant; **: p < 0.01; ***: p < 0.005; ****: p < 0.001. **D**, ΔT_vac lys_. CTRL: 35 foci, N = 7; Blebbistatin: 34 foci, N = 7; SCRAMBLE: 35 foci, N = 7; MyoIIAB siRNA: 34 foci, N = 7. **E**, actin foci duration. CTRL: 17 foci, N = 4; Blebbistatin: 30 foci, N = 3; SCRAMBLE: 17 foci, N = 4; MyoIIAB siRNA: 21foci, N = 4. **F**, percent of SCV with actin cups. SCRAMBLE: 27 SCVs, N = 3. MyoIIAB siRNA: 34 SCVs, N = 5. Blebbistatin: 42 SCVs, N = 7. Chi-square. *: p = 0.031. **G**, Scheme of myosin II activation and putative role in SCV rupture. Local Ca^2+^ increase (pale green) lead to myosin II activation (green circles) in membrane ruffles induced by invading *Shigella* (blue). Activated myosin II in ruffles or associated with the retrograde flow (purple arrow) exerts pulling forces on the SCV membrane (dark pink) transduced by actin filaments (light pink) and the actin cup / coat (dark pink).

Treatment with the myosin II inhibitor blebbistatin or anti-myosin II siRNA led to a two-fold increase in the ΔT_vac lys_, consistent with a role for myosin II in vacuolar rupture (Fig. 4D). This delay in vacuolar rupture was also associated with an increased duration of actin polymerization at *Shigella* invasion sites combined with a decrease in SCVs associated with actin cups, mirroring the effects observed in the absence of extracellular Ca^2+^ (Figs. 4E, F and 3D, E).

These results suggest a role for myosin II and actomyosin contraction in the rupture of the SCV. The transient up-regulation of myosin II at the onset of *Shigella* invasion correlates with the peak of long lasting local Ca^2+^ increases elicited at bacterial entry sites during the first five minutes following bacterial challenge. We suggest that myosin II exerts contractile forces transduced by actin filaments surrounding the vacuole thereby contributing to SCV rupture while it is activated at actin projections at a distance from the bacterium within its SCV (Fig. 4G).

### Ca**^2+^**-dependent recruitment of myosin VI at the SCV precedes vacuolar rupture

Among non-conventional myosins, myosin VI is an actin minus-end motor involved in generating tension and membrane curvature at the plasma membrane during endocytosis, secretion and autophagy (Houdusse and Sweeney, 2007; Tomatis et al., 2013; de Jonge et al., 2019). Myosin VI is directly regulated by Ca^2+^ calmodulin (Batters et al., 2016), unlike Myosin II whose activation requires a Ca^2+^-sensitive kinase. Ca^2+^ calmodulin up-regulates the recruitment of myosin VI by relieving auto-inhibition, allowing to switch on the motor promoting force production (Altman et al., 2004; Batters et al., 2016). Once activated and recruited, the Myosin VI motor can perform distinct function depending on the force it is working under. Specifically, Myosin VI can tether actin structures to membranes and may either anchor the SCV to the actin cortex, play a role of a transporter or an actin organizer (Frank et al., 2004; Loubery and Coudrier, 2008). Previous *in vitro* experiments indicated in fact that the processivity of the dimeric Myosin VI motor was lost in the presence of Ca^2+^, which may thus lead to reduced efficiency as a transporter (Yoshimura et al., 2001; Morris et al., 2003). Myosin VI therefore represents a good candidate to anchor the SCV at the actin cortex in a Ca^2+^-dependent manner, thereby facilitating vacuolar membrane rupture upon the exertion of mechanical forces.

We next investigated the localization of Myosin VI during rupture of the SCV by performing live imaging of cells transfected with GFP-myosin VI and Gal3-mOrange. As illustrated in Fig. 5A and as previously reported, GFP-myosin VI labeled motile vesicles consistent with its role in endocytosis (Buss et al., 2001; Engevik and Engevik, 2022). Strikingly, GFP-myosin VI also labeled distinct motile patches that clustered at *Shigella* invasion sites and transiently interacted with the SCV (Fig. 5A, arrows; Movie EV4). Quantitative analysis of GFP-myosin VI fluorescence intensity indicated that the peak of its recruitment at invasion sites preceded vacuolar rupture (Figs. 5B, C, arrowheads and red arrows). In some instances myosin VI recruitment could occur several minutes before vacuolar rupture (Fig. 5B, blue and green traces), but in the large majority of events, it only preceded vacuolar rupture by a few seconds (Fig. 5B, red traces and Fig. 5C), with an average time duration of 4.7 seconds, as estimated from the intersection of the linear fit with the x-axis when comparing the time corresponding to the peak of myosin VI recruitment to that of ΔT_vac lys_ (Fig. 5C).

**Figure 5.**
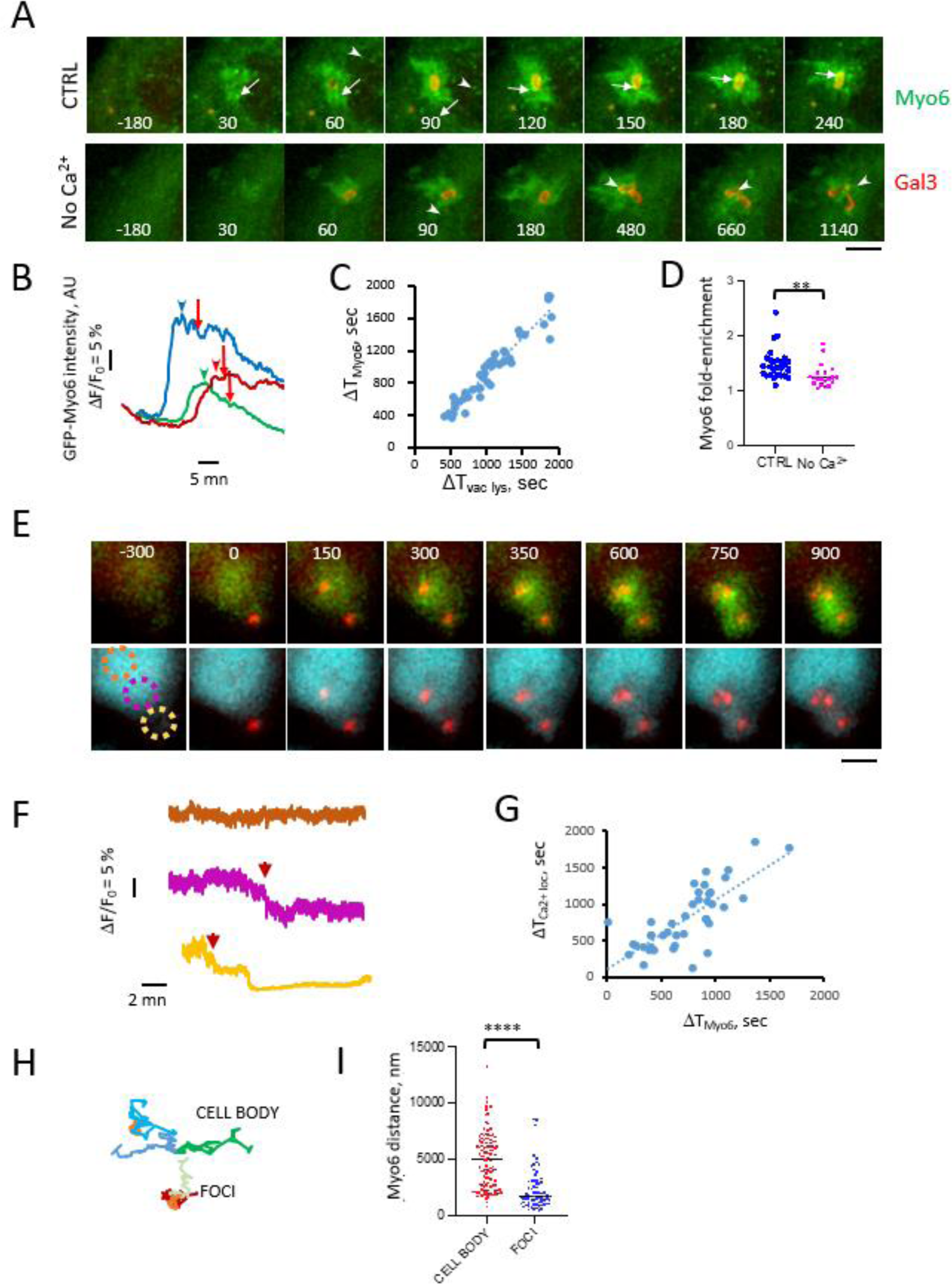
Myosin VI recruitment at the SCV is associated with long lasting local Ca^2+^ increases and precedes vacuolar rupture. Gal3-mOrange and GFP-Myosin VI double-transfected cells were challenged with *Shigella* in the presence (CTRL) or absence (No Ca^2+^) of external Ca^2+^. Samples were imaged for GFP-Myosin VI and Gal3-mOrange fluorescence by spinning disk confocal microscopy. **A**, Representative time-series of micrographs corresponding to maximal z-projection. Numbers: time relative to the start of actin foci formation in min. Arrows: large GFP-Myosin VI clusters at *Shigella* invasion sites. Arrowheads: smaller GFP-myosin VI patches elsewhere in the cell, or in foci in absence of external Ca^2+^. Red: Gal3-mOrange; Green: GFP-myosin VI. Scale bar = 5 μm. **B**, representative traces of the average GFP-myosin VI average fluorescence intensity at bacterial invasion site over time. Arrowheads: peak of GFP-myosin VI intensity. Red arrows: SCV rupture. **C**, Time of myosin VI recruitment peak plotted as a function of ΔT_vac lys_. Pearson correlation coefficient of the linear fit = 0.91. **D**, GFP-myosin VI fold-enrichment at *Shigella* invasion sites. CTRL: 26 foci, N = 5. No Ca^2+^: 20 foci, N = 4. **E-G**, Gal3-mOrange and GFP-myosin VI double-transfected cells were loaded with Fura 2-AM, challenged with *Shigella*, and subjected to Ca^2+^ and GFP-myosin VI imaging (Methods). **E**, representative time-series of fluorescence micrographs. Numbers: time relative to the start of actin foci formation in seconds. Scale bar = 5 μm. Top panels: maximal z-projection of Gal3-mOrange (red) and GFP-myosin VI (green) planes. Bottom panels: Fura-2 (cyan) and Gal3-mOrange (red) fluorescence. **F**, traces corresponding to variations in intracellular Ca^2+^ in the cell region in depicted in panel E with matching color. **G**, ΔT_Ca_^2+^ _loc_ plotted as a function of ΔT_Myo6_. Pearson correlation coefficient of the linear fit: 0.58. **H**-**J**, GFP-myosin VI clusters or patches as shown in panel A, were tracked over time at *Shigella* invasion sites (FOCI) or in the rest of the cell body (CELL BODY). **H**, representative tracks. **I**, **J**, GFP-myoVI patches or clusters’ distance covered (**I**) and velocity (**J**) in *Shigella* invasion sites or in the cell body. 70 foci clusters, N = 3. 98 cell body patches, N = 10.

Myosin VI clustering and recruitment at *Shigella* invasion sites was not observed in the absence of extracellular Ca^2+^, suggesting a role for Ca^2+^ influx and long lasting local Ca^2+^ increases (Figs. 5A, D, No Ca^2+^; Movie EV5). To further analyze the association between these long lasting local Ca^2+^ increases and myosin VI recruitment during vacuolar rupture, we performed dual live fluorescence imaging of GFP-myosin VI and Ca^2+^. As illustrated in Figs. 5E, F, GFP-myosin VI recruitment at the SCV spatio-temporally correlated with long lasting local Ca^2+^ increases as well as vacuolar rupture. Consistently, a linear correlation was observed between the peak recruitment of myosin VI at the SCV and the elicitation of these long lasting local Ca^2+^ increases (Fig. 5G).

Because long lasting local Ca^2+^ increases occurred at *Shigella* invasion sites, we expected GFP-myosin VI patches to be less motile at these sites relative to patches in other areas of the cell cytosol, if Ca^2+^ was triggering motor recruitment. Indeed, and as illustrated in Fig. 5H, tracking analysis indicated that GFP-myosin VI patches remained localized at the invasion sites for extended duration with an average traveled distance over the acquisition period that was 1.8 fold less than that of patches localized in other areas of the cytosol (Fig. 5I).

Together, these results indicate that myosin VI is recruited at *Shigella* invasion sites in a Ca^2+^-dependent manner, and that its recruitment is associated with long lasting local Ca^2+^ increases linked to vacuolar rupture. The decreased motility of GFP-MyoVI patches in invasion foci is consistent with their restricted diffusion promoted by local Ca^2+^ increase and their association with SCVs.

### Myosin VI tethers the SCV to the cell cortex

We next investigated the role of myosin VI in rupture of the SCV. As shown in Fig. 6A, cells treated with anti-myosin VI siRNA showed a 2.5 increase in ΔT_vac lys_ relative to control cells (Fig. 6A, compare siRNA MyoVI and SCRAMBLE). A similar increase in ΔT_vac lys_ was observed when cells were treated with the myosin VI inhibitor 2, 4, 6-triiodophenol (TIP) (Heissler et al., 2012; Fig. 6A, compare TIP and CTRL).

**Figure 6.**
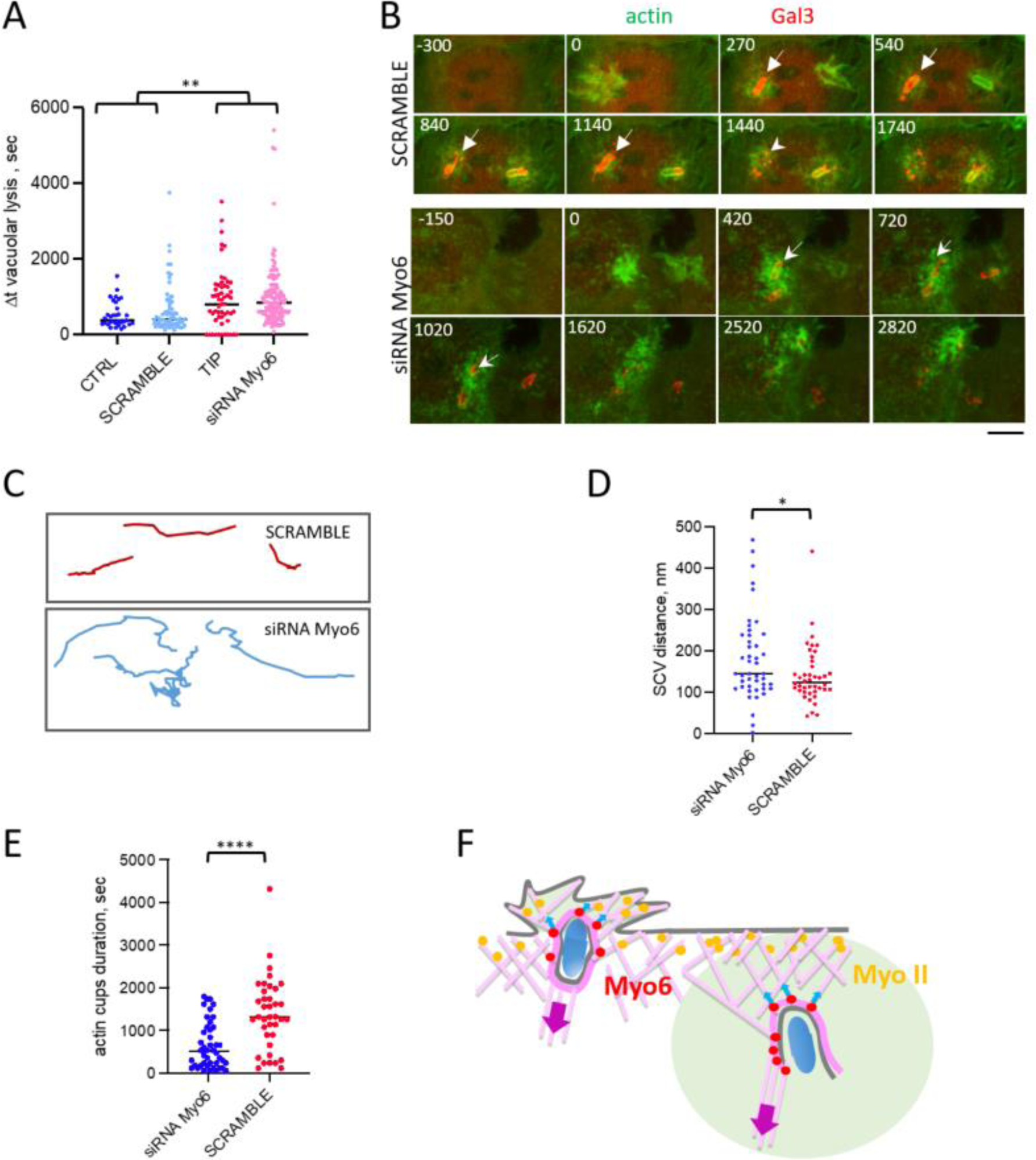
Myosin VI tethers the SCV to the cell cortex. Cells double-transfected with Gal3-mOrange and GFP-actin were challenged with *Shigella* and processed for live fluorescence imaging. **A**, ΔT_vac lys_ determined for the indicated samples. Cells treated with: 25 μM of the myosin VI inhibitor (TIP), 35 SCVs, N = 4; medium alone (CTRL), 28 SCVs, N = 4; scramble siRNA control (SCRAMBLE), 28 SCVs, N = 4); anti-myosin VI siRNA (siRNA MyoVI), 69 SCVs, N = 5. **B**, Representative time-series of micrographs corresponding to maximal z-projections. Numbers: time relative to the start of actin foci formation in seconds. Red: Gal3-mOrange; Green: GFP-actin. Scale bar = 5 μm. The arrows point at a fixed position in the field. Note the following rupture, the SCV remains immobile in cells treated with scramble but not anti-myosin VI siRNA. **C**-**E**, SCV were tracked for the first 10 min immediately following rupture (Methods). **C**, representative tracks. **D**, SCV velocity in nm / sec. **E**, distance covered by the SCV. SCRAMBLE: 42 SCVs, N = 5. siRNA MyoVI: 43 SCVs, N = 6. **F**, actin cup duration. SCRAMBLE: 38 SCVs, N = 5. siRNA MyoVI: 43 SCVs, N = 6. Mann-Whitney test. **: p < 0.01; ****: p < 0.001. **G**, Scheme of Ca^2+^ regulation of myosin II and myosin VI during SCV rupture. Local Ca^2+^ increase (pale green) trigger the tethering of myosin VI (red circles) to the SCV (dark pink) and the transduction of forces (blue arrows) via actin filaments (light pink) exerted by myosin II (yellow circles) and the retrograde flow (purple arrow). The combination of myosin II-based pulling forces and constraining of SCV in the actin cortex by myosin VI favors SCV rupture.

The delay in vacuolar rupture associated with myosin VI inhibition correlated with a striking higher tumbling motility of SCV membranes in the cell cortex and as well as a lesser integrity of SCV membranes following rupture. In control conditions, Gal3-labeled SCV remnants were observed to preserve their shape for several minutes, being either non motile or moving several micrometers towards the cell interior in an anisotropic manner along the retrograde flow (Figs. 6B, SCRAMBLE, arrows; Figs 6C-E, SCRAMBLE; Movies EV6 and EV7). Under these conditions, SCVs eventually lost their integrity and disintegrated before disappearing (Ehsani et al., 2012; Chang et al., 2024; Fig. 6B, SCRAMBLE, arrowhead; Movies EV6 and EV7). In contrast, in anti-myosin VI siRNA-treated cells, SCVs showed a higher degree of tumbling motility following rupture (Fig. 6B, siRNA MyoVI, arrows; Movies EV8 and EV9), or traveling over longer distances and in less anisotropic manner than in control cells (Figs. 6C, D. compare SCRAMBLE and siRNA MyoVI).

When analyzing the dynamics of actin structures, as observed for the depletion of external Ca^2+^ and myosin II inhibition, we found that myosin VI inhibition destabilized the actin cups / cocoon forming around the SCV. These structures disintegrated 2.1 faster than in control cells, with an average duration time ± SEM of 10.7 ± 1.4 min compared to 23.0 ± 2.3 min for control cells (Fig. 6B, siRNA MyoVI, arrowhead; Figs 6C-E, siRNA MyoVI; Movies EV8 and EV9).

These results indicate that myosin VI is involved in rupture of the SCV and constrains its motility in the cell cortex. Together, our findings suggest that long lasting local Ca^2+^ increases allows myosin VI to tether the SCV to the cortical actin network, favoring the formation of the associated actin cup / cocoon and constraining its motility in the cell cortex. This myosin VI-dependent tethering enables mechanical forces exerted on the SCV to efficiently rupture it (Fig. 6F).

## Discussion

Despite their role in infectious pathology, the disassembly of vacuoles containing intracellular bacterial pathogens is a process that has been only recently been investigated with the aim to decipher the underlying molecular mechanisms (Pizarro-Cerda et al., 2016; Gutierrez and Enninga, 2022). In the case of *Shigella*, components of recycling and exocytic compartments are recruited at macropinosomes contacting the SCV, but these rather appear to contribute to the dispersal of the SCV membranes following rupture. The host dynein motor protein complex, possibly recruited through the macropinosomes, has been shown for a full unwrapping of the damaged SCV remnants (Chang et al., 2024). Together, these works have determined that rupture of the SCV during *Shigella* invasion takes place through successive steps with an early damage of the vacuole and a later unpeeling of the broken membrane remnants. How early SCV rupture occurs though has remained unclear.

Actin-based forces have been implicated in the breakdown of the envelope of the Starfish oocyte large nuclei during mitosis (Mori et al., 2014). In this model, nuclear envelope breakdown (NEBD) is facilitated and preceded by the formation of a dense actin structure or actin shell that line the nuclear inner membrane, from which Arp2/3-dependent actin spikes were observed to protrude into the nucleus and proposed to pierce the inner and outer nuclear membranes (Mori et al., 2014). At a different scale, in the case of the SCV, this actin shell is reminiscent of the actin cups and cocoon forming around the SCV. Similar to the nuclear actin shell, the SCV actin cups and cocoon have been found to facilitate its rupture, although these latter form on the cytosolic side of the vacuole while the actin shell form inside the nucleus (Kuhn et al., 2020). As for NEBD, actin is not essential for the rupture of the SCV, but it accelerates the process suggesting the implication of alternate mechanisms, possibly involving the T3SS for the initial damage and microtubules for the full unwrapping of the SCV membranes. Timing represents a critical factor during mitosis, where a rapid NEBD is required for proper chromosomal segregation. Rapid escape from the vacuole could be also critical for *Shigella* to evade detection and cellular immunity mechanisms (Torraca and Mostowy, 2016; Chang et al., 2024). The common involvement of an actin structure suggests that tethering of membranes to a supporting scaffold facilitates its rupture, perhaps by constraining the membranes where local tearing forces are exerted.

Actin can generate protruding forces via its polymerization or pulling forces via myosin motors. *Shigella* invasion involves type III effectors that trigger actin polymerization and actomyosin contraction (Valencia-Gallardo et al., 2015). Since the majority of events of vacuolar rupture occurs in the continuation of the invasion processes, such forces are likely to be exerted at the SCV membranes. The specific organization of these forces and their potential implication, however, need to be considered in the context of the actin dynamics during the invasion process. *Shigella*-induced actin polymerization accounts for ruffles forming around invading bacteria, while actomyosin contraction is expected to pull the bacteria in the cell (Fig. 6F). The deep mechanism underlying the spatio-temporal regulation and concerted activity of the responsible type III effectors is not fully understood, even though it has been found that one effector, IcsB reprograms host GTPases and alters the formation of the actin cocoon (Liu et al., 2018; Kühn et al., 2020). IpaA, another injected *Shigella* type III effector, targets vinculin and talin to cap actin filaments and prevent actin polymerization at the bacteria-cell contact site, contributing to the formation of the initial actin cups anchoring bacteria to the cytoskeleton within the cortex to allow their pulling inside the cell (Fig. 6F; Valencia-Gallardo et al., 2019; Valencia-Gallardo et al., 2023).

Once bacteria are internalized into a SCV, polymerization of uncapped actin filaments may exert forces at the SCV membrane resolved from plasma membrane ruffles (Fig. 6F). Upon vacuolar rupture, these forces linked to actin polymerization may be visualized by the propelling of SCV remnants that we observed (Movie EV6). Consistently, the actin comets exhibit long trajectories suggesting their association with small vacuolar membrane debris while comet tails undergoing short and swirling or circular trajectories may be associated with membrane remnants still connected to large damaged parts of the SCV, as being held by a “lasso” (Movie EV7). These actin-polymerization based forces do not appear to depend on Ca^2+^ influx, and propelling of SCV remnants by actin comets was even exacerbated in the absence of external Ca^2+^. It is possible that these Ca^2+^ independent events require the post-translational reprogramming of actin-modulators directly at the SCV, for example through the bacterial acylase IcsB (Liu et al., 2018).

In contrast, we show that local Ca^2+^ increases regulate the activation of myosin II at membrane ruffles surrounding the forming SCV. We found that myosin II is involved in SCV rupture. Blebbistatin that specifically inhibits the motor activity of myosin II delays vacuolar rupture consistent with actomyosin based forces contributing to the tearing of SCV membranes. Our findings also indicate that myosin II antagonizes actin polymerization in ruffles surrounding the forming SCV while stabilizing actin cups / cocoons. The antagonistic actin polymerization / actomyosin contraction activities is well documented during cell adhesion (Shaks et al., 2019). During cell adhesion, Arp2/3-dependent actin polymerization downstream of the small GTPase Rac allows lamellae extension and formation of focal complexes, whereas actomyosin contraction downstream of the Rho GTPase favors the maturation of focal complexes into bona fide focal adhesions in a substrate stiffness-dependent manner (Shaks et al., 2019). During mechanotransduction, the switch between actin polymerization and actomyosin contraction strengthens adhesion structures. Upon invasion of bacteria such as *Shigella*, counterforces are not expected to be sufficient to trigger the maturation of focal complexes into focal adhesions. However, the IpaA type III effector tricks cells into forming a pseudo-focal adhesion at the bacterial contact site, which may contribute to the transduction of actomyosin-based forces during and following invasion and at SCV membranes. Along these lines, the myosin II-dependent strengthening of actin cups and cocoon surrounding the SCV may show similarities with the actomyosin-dependent strengthening of adhesion structures associated with increased pulling forces (Sun et al., 2016). The presence of phospho-myosin in membrane ruffles associated with bacterial invasion and surrounding the SCV suggests that myosin II could also exert contractile forces on the SCV via actin filaments in these ruffles independent of those exerted by the retrograde flow (Fig. 6F). The link between actin polymerization in ruffles and formation of an actin cocoon around the SCV suggests that these contractile forces involve the tethering of actin filaments at the plasma membrane, perhaps combined with the tension of the plasma membrane in ruffles as proposed at a larger scale for the cycle of extension and retraction of the lamella (Ryan et al., 2017).

For pulling forces to be effective in tearing on the SCV membrane, this latter needs to be constrained. We found that long lasting Ca^2+^ increases trigger the recruitment and tethering of myosin VI patches at the SCV. As observed for Ca^2+^ and myosin II, myosin VI stabilize the actin cup / cocoon juxtaposing the SCV, with an increased integrity of the SCV. We also found that myosin VI is required for efficient vacuolar rupture. Additionally, we found that myosin VI was recruited to *Shigella*-induced membrane ruffles, similar to other studies showing myosin VI recruitment to epidermal growth factor-induced ruffles (Chibalina et al., 2009; Buss et al., 1998). In line with its various functions at different subcellular location, myosin VI interacts with diverse molecular partners (Magistrati and Polo, 2021). Myosin VI is also found at the leading edge of Drosophila ovary border cells, in complex with E-cadherin and beta-catenin, where it was proposed to contribute to cell motility by pushing actin filaments outwards via its minus-end motor activity (Geisbrecht and Montell, 2002; Breshears and Titus, 2007).

Myosin VI was also observed in ruffles induced by enteroinvasive *Salmonella*, where together with the bacterial effector SopB, a phosphatidylinositol polyphosphate phosphatase / isomerase, it contributes to the accumulation of PI(3)P, PI(3,4)P2, and PI(3,4,5)P3 at invasion foci. Indeed, depletion of myosin VI results in a defect in ruffle formation and bacterial invasion (Brooks et al., 2017). The production of PI(3)P at the SCV by SopB also requires the recruitment of Rab5 and the activation of the class III PI3K (Mallo et al., 2008). SopB also helps fusion of the *Salmonella* vacuole with surrounding macropinosomes allowing its growth for vacuolar stabilization (Stévenin et al., 2019). In light of more recent work, defects in ruffle formation in myosin VI-depleted cells may be linked to the mislocalization of Rab5-positive endosomes activating the PI3K-Akt pathway (Masters et al., 2017). Interestingly, despite the fact that *Shigella* expresses IpgB1 and IpgD, the respective type III effectors orthologs of SopE and SopB, we did not find evidence for a role of myosin VI in *Shigella*-induced actin ruffles or bacterial invasion (Fig. EV4). Also, *Salmonella* does not form large actin cups or cocoons and its vacuole is not held within the cellular cortex as the *Shigella* vacuole. Interestingly, this lack of tethering to the bacterial vacuole to the cortex in the case of *Salmonella* coincides with inefficient vacuolar damage.

Our results indicate that myosin VI-based forces contribute to the efficiency but are not essential for vacuolar rupture at the SCV. Strikingly, we found that local Ca^2+^ increases coincided with the formation and tethering of myosin VI patches at the SCV, preceding the rupture event. By tracking SCVs, we found that their degree of motility is largely increased upon myosin VI inhibition consistent with its role in constraining the SCV in the actin cortex. In myosin VI depleted cells, the delay in vacuolar rupture was associated with less vacuole stability and less constraints led to more mobile SCV remnants. These findings are consistent with the regulatory role of Ca^2+^ on myosin VI inferred from structural and *in vitro* gliding actin filament studies (Yoshimura et al., 2001; Morris et al 2003; Batters et al., 2016). In these studies, Ca^2+^ was shown to promote conformational changes in myosin VI by regulating the binding of calmodulin to different sites, leading to joint increase in lipid cargo binding and destabilization of the lever arm leading to lack of motility (Batters et al., 2016).

To the best of our knowledge, our studies are first to support a role for local Ca^2+^ increases in tethering intracellular membranes to the actin cortex by myosin VI. The tethering role of myosin VI was evocated for signaling endosomes driving Akt activation, since depletion of myosin VI by siRNA led to a shift from the cell cortex to a perinuclear location (Masters et al., 2017). Further investigating will detail which partners on the SCV can lead to myosin VI recruitment dependent on Ca^2+^ activation, thus leading to the transition of myosin VI from a location on endosomes to patches contacting the SCV. We propose that upon local Ca^2+^ increase, myosin VI-based tethering of the SCV to the actin cortex, acting as a “holding hand”, facilitates the early steps of its rupture by force exerted by the myosin II-based contraction acting as the “pulling hand”. Such function could also be relevant for the role myosin VI plays in trafficking processes involving local Ca^2+^ influx and actin at the plasma membrane, such as endocytosis, macropinocytosis or exocytosis.

## Methods

### Antibodies and reagents

The anti-*Shigella* serotype V LPS antibody was described previously (Valencia-Gallardo et al., 2019). Rabbit polyclonal antibodies against phospho-myosin light chain 2 (Ser219) (#3671), myosin IIA (#3403), myosin IIB (#8824), myosin VI (#13592) were from Cell Signaling. Secondary anti-rabbit IgG conjugated to Alexa Fluor R 488 (#111-545-003), Alexa Fluor R 594 (#111-585-003), Alexa Fluor R 647 (#111-605-003), or horseradish peroxidase (#111-035-003) and anti-mouse IgG conjugated to Alexa Fluor R 488 (#115-545-003) horseradish peroxidase (#115-035-003) were from Jackson Immunoresearch. Alexa Fluor 647 -conjugated Phalloidin (#A22287) was from Invitrogen. The anti-STIM-1 mouse monoclonal antibody clone 5A2 (#WH0006786-M1), Blebbistatin (#203391), the myosin VI inhibitor 2, 4,6-triiodophenol (TIP) (#137723) were from Sigma-Aldrich. Fluo-4, AM (#F14201), Fura-2, AM (#F1225) and the anti-beta-actin BA3R (#MAS-15739) antibody was from ThermoFisher Scientific.

### Cell lines, bacterial strains and culture conditions

HeLa cells (ATCC CCL-2) were grown in DMEM medium supplemented with 10% fetal bovine serum (FBS) in a 37C incubator with 10 % CO_2_. *Shigella* wild-type serotype V M90T and its isogenic non-invasive *mxiD* mutant were described previously (ref). Bacterial strains were grown in trypticase soy broth at 37C. Cells were challenged with bacteria, as previously described (ref). Briefly, bacteria were grown to an OD_600 nm_ of 0.6–0.8, washed three-times by successive centrifugation at 13 kg for 30 s and resuspended in EM buffer (120 mM NaCl, 7 mM KCl, 1.8 mM CaCl_2_, 0.8 mM MgCl_2_, 5 mM glucose, and 25 mM HEPES, pH = 7.3) at a final OD_600 nm_ of 0.1. For analysis of fixed samples, cells were incubated with the bacterial suspension for 15 min at 21°C prior to incubation at 37°C for the indicated time period. Samples were fixed with PBS containing 3.7% PFA for at 30-60 min before processing for immunofluorescence microscopy. For live sample imaging, the coverslip was mounted in a microscopy chamber and bacterial challenge was performed on a 37°C-heated microscope stage. When indicated, blebbistatin and TIP, at a final concentration of 50 μM and 5 μM, respectively, was added 30 minutes at 37°C prior to bacterial challenge.

### Plasmid constructs, siRNAs, and cell transfection

The pEGFP-myosin VI expressing full-length myosin VI fused to the C-terminus of GFP was previously described (Warner et al., 2003). The previously described siRNAs were used: hSTIM1-1140 siRNA (sense, 5′GGCUCUGGAUACAGUGCUCtt3′, antisense, 5′-GAGCACUGUAUCCAGAGCCtt3′). The scrambled dsRNA (sense, 5′GUGCGACUGCUGGACUACUtt3′, antisense, 5′AGUAGUCCAGCAGUCGCACtt3′) was used as a negative control (Jousset et al., 2007; (Dharmacon, ON-TARGETplus Mouse *Orai1* siRNA, L-056431-02-0005). Stealth siRNAs set of HSS106852, HSS106853, HSS106854 and set of HSS106897, HSS106898, HSS106899 (ThermoFisher Scientific) were used for knock down of myosin IIA, B and VI, respectively.

### Cell transfection

For transfection experiments, cells were seeded at 2.5 x 10^4^ cells on 28 mm-diameter coverslips in a 6-well plates 48 hours before the experiment. After 16 h, cells were transfected with 0.5 to 1.0 μg of plasmid using 6 μL JetPEI transfection reagent (Polyplus) for 16 h following the manufacturer’s recommendations. For siRNA transfections, cells were transfected with 1.5 μl of 20 μM siRNA using 9 μl of lipofectamine.

### Immunofluorescence analysis

Fixed samples were permeabilized by incubation in PBS containing 0.1% Triton X-100 for 4 minutes at 21C. Samples were washed three times with PBS and blocked in PBS containing 1% FBS for 30 minutes, prior to incubation with the primary antibody at the following dilution anti-LPS (1 :5000), anti-STIM-1 (1:2000), anti-phosphomyosin LC2 (1:200) in PBS containing 0.2% BSA, followed by incubation with secondary anti-IgG antibody and Alexa-Fluor phalloidin at 1:200 dilution. Samples were analyzed using an Eclipse Ti inverted microscope (Nikon) equipped with a 60×objective, a CSU-×1 spinning disk confocal head (Yokogawa), and a Coolsnap HQ2 camera (Roper Scientific Instruments), controlled by the Metamorph 7.7 software. Analysis of fluorescent actin filaments was performed using a Leica confocal SP8 using a 63× objective.

### Live Dual fluorescence imaging of vacuolar rupture

For dual Ca^2+^ and SCV rupture imaging, cells transfected with Gal3-mOrange were loaded with 3 μM Fluo-4, AM or Fura-2, AM in EM buffer as previously described (Sun et al., 2017). Samples were mounted on a microscopy chamber on a 37°C-heated stage on an inverted Leica DMRIBe fluorescence microscope, equipped with LED illuminating sources and a Cascade 512B EM-CCD back-illuminated camera driven by the Metamorph software 7.7 from Roper Scientific Instruments. Acquisition was performed using a 63/1.25 HCX PL APO objective and fluorescence filters (Fura-2: excitation BP360 ± 40 nm, emission BP470 ± 40 nm; Fluo-4: excitation 480 ± 20 nm, emission 527 ± 30 nm; Gal3-mOrange: excitation BP540 ± 20 nm, emission LP590 nm). Time points were acquired every 3 seconds for a whole duration of up to 90 min. For Fluo-4 and Fura-2, one image corresponding to the focus plane was acquired for each time point. For Gal3-mOrange, a stack of 5-10 Z-planes spaced by 400 nm were acquired in stream mode to cover the cell volume. For dual imaging of GFP-actin or GFP-myosin VI and Gal3-mOrange, samples were analyzed using an Eclipse Ti inverted microscope (Nikon) equipped with a Plan Apo lamD 60x (NA 1.42) objective, a CSU-×1 spinning disk confocal head (Yokogawa), a sCMOS 2034 camera (Hamamatsu), 488 nm and 561 nm 100 mW lasers, using a BP525 ± 60 nm and BP600 ± 52 nm emission filters, controlled by the Metamorph 7.7 software. Time points were acquired every 15 seconds or 30 seconds, with 24-30 Z-planes spaced by 400 nm in stream mode for each wavelength.

### Image analysis

Changes in the Fluo-4 or Fura-2 fluorescence intensity (ΔF) were calculated relative to the resting ratio or fluorescence value (F_0_) as ΔF/F_0_. The percent of cells showing global or local Ca^2+^ increase at least three-fold above background noise during the length of the analysis was determined. All images were corrected for background fluorescence. The start of foci formation (T_0_) was determined as the inflexion point between resting and increased average fluorescence intensity levels in the region corresponding to the invasion site. ΔT_vac lys_ was determined as the difference between the time point corresponding to vacuolar rupture based on Gal3 labeling of the SCV and T_0_. ΔT_Ca2+ glob_ and ΔT_Ca2+ loc_ was determined as the time interval between the peak of global and local Ca2+ increase relative to T_0_, respectively.

The analysis of STIM-1, phospho-myosin LC II, GFP-actin and GFP-myosin VI recruitment at *Shigella* invasion sites was performed on average intensity of confocal planes Z-projections of immunofluorescent labeled samples as previously described (Tran Van Nhieu et al., 2013). The fold-enrichment at invasion site was determined as the ratio of the average fluorescence intensity corrected for background in the invasion site outlined by the actin staining over that of the remaining cell body. The peak of GFP-Myosin VI corresponded to the time point with the highest average intensity value preceding vacuolar rupture.

The duration of actin foci formation was calculated as the time period between the inflexion points corresponding to increase in average fluorescence intensity and return to baseline levels. Invasion site were scored positive for the presence of an actin cup when at least half of the SCV or forming SCV was associated with an F-actin coat.

Tracking of GFP-myosin VI patches and clusters, and of the Gal3-mOrange labeled SCV was performed using the Metamorph 7.7 tracking module. SCV tracking was performed for a 300 seconds-time period immediately following rupture to avoid variations linked to subsequent disintegration of SCV membranes.

### Statistics

Actin coat-structure formation was analyzed using in a Pearson Chi-square test (R Statistical Software). For all statistical tests a Shapiro-Wilk normality test was performed to apply a parametric or a non-parametric test. The statistical test used, number of data points (n) and number of experimental replicates (N) are indicated in the figure legend.

## Acknowledgements

We thank Tristan Piolot from the CIRB imaging facility for technical help. This work was funded by the Inserm, the CNRS and the Collège de France, as well the ANR grant PureRuptureMag and RabReprogram. MB was a FRM (Fondation pour la Recherche Médicale) post-doctoral fellowship recipient. CHS was funded by the Chinese Science Council. JE is member of the LabEx IBEID and Milieu Interieur, and he acknowledges support by the EU (ERC-CoG EndoSubvert).

## Author contributions

M.B. designed, performed, analyzed experiments and wrote the manuscript. CH.S. performed and analyzed experiments. J.E. and A.H. analyzed data and wrote the manuscript. GTVN supervised the work, analyzed data and wrote the manuscript.

---

**Figure EV1.**
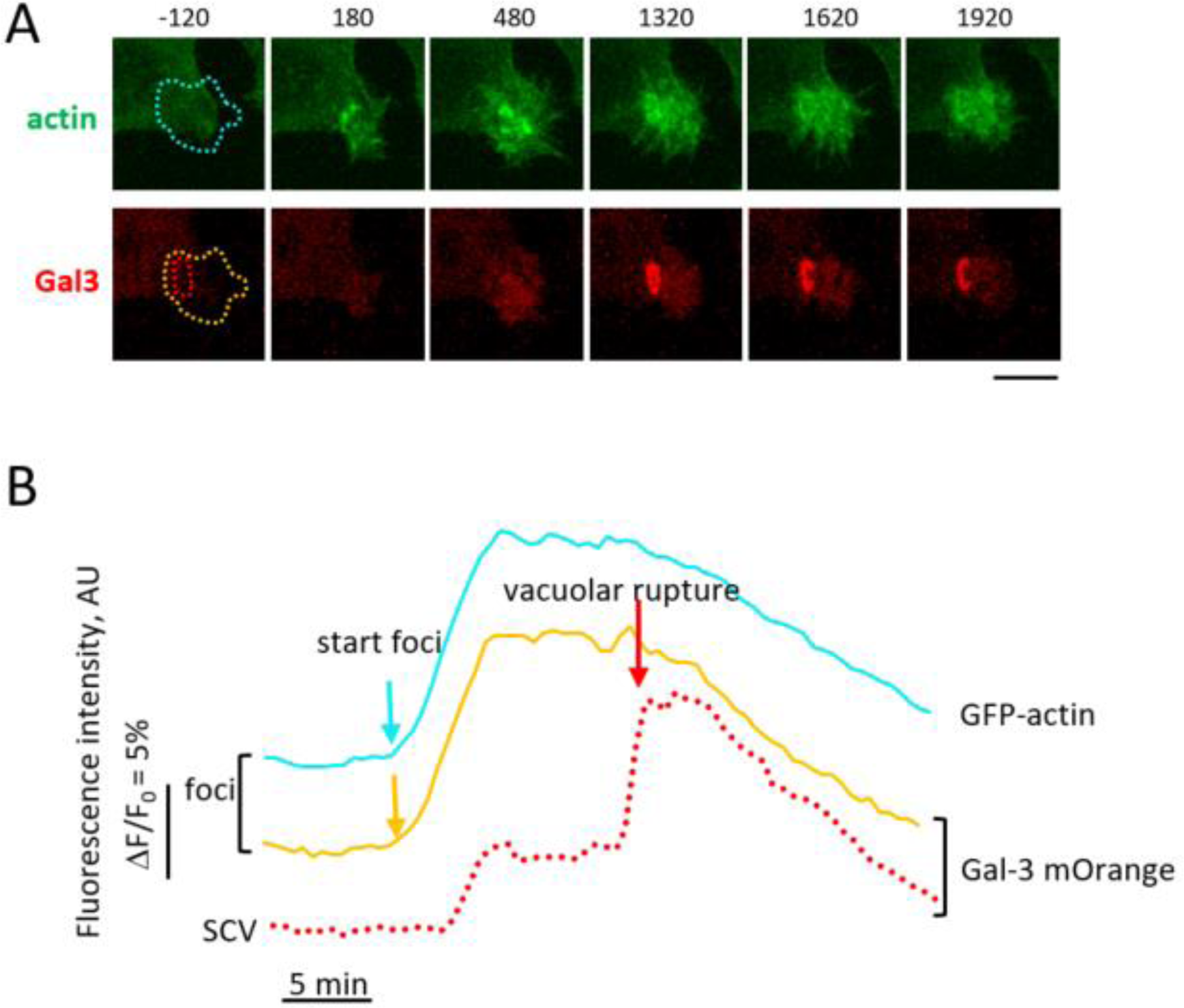
Gal3-mOrange recruitment as a proxy for foci formation. Cells double-transfected with Gal3-mOrange and GFP-actin were challenged with *Shigella* and processed for live fluorescence imaging. **A**, representative time-series of micrographs corresponding to maximal z-projections. Numbers: time relative to the start of actin foci formation in seconds. Red: Gal3-mOrange; Green: GFP-actin. Scale bar = 5 μm. **B**, average fluorescence intensity of the regions depicted in A with matching color, corresponding to GFP-actin (cyan), Gal3-mOrange in foci (yellow), or Gal3-mOrange at the SCV (dotted red).

**Figure EV2.**
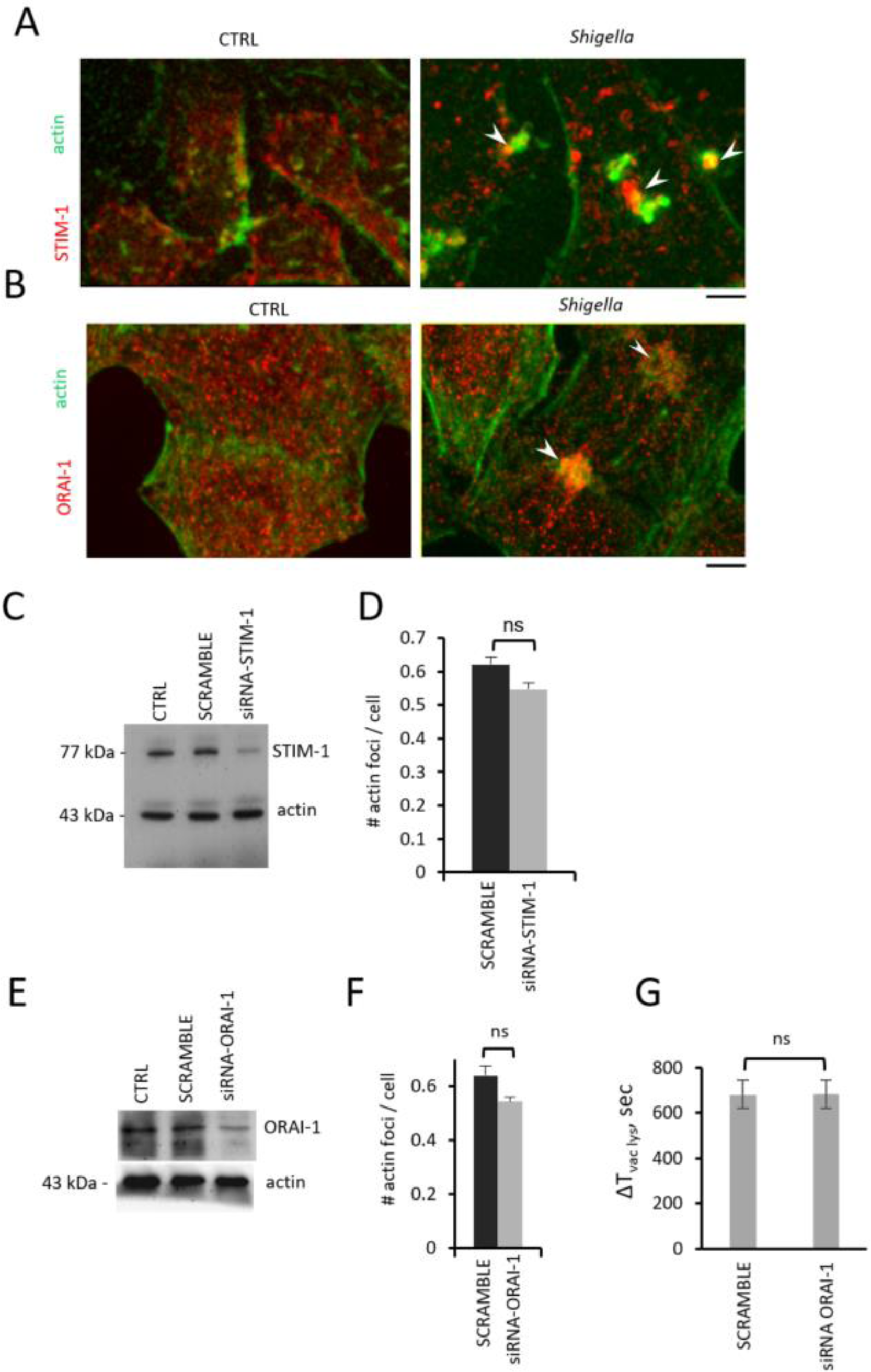
Effects of siRNA-mediated depletion of STIM-1 and ORAI-1. A, B, Representative immunofluorescence micrographs. CTRL: uninfected cells. *Shigella*: cells infected for *Shigella* for 15 min at 37°C. Staining: green: F-actin; red: STIM-1 (A) or ORAI-1 (B). Arrows: STIM-1 recruitment at invasion sites. Arrowheads: ORAI-1 recruitment at invasion sites. Scale bar 5 μm. **C**-**F**, cells were transfected with anti-STIM1 (**C**, **D**) or anti-ORAI-1 (**E**-**F**) siRNAs. **C**, **E**. Cell lysates were analyzed by Western-blot following SDS-PAGE in a 10 % polyacrylamide gel using anti-STIM-1 (**C**), anti-ORAI-1 (**E**), or anti-actin antibody as a loading control (**C**, **E**). **D**, **F**, cells were challenged with wild-type *Shigella*, fixed and processed for fluorescence staining. The number of *Shigella* actin foci per cell were scored by fluorescence microscopy analysis for the indicated samples. **D**, SCRAMBLE: 1487 cells, 865 foci, N = 3. siRNA STIM-1: 1884 cells, 1116 actin foci. N = 3. **F**, SCRAMBLE: 1163 cells, 658 foci, N = 3. siRNA ORAI-1: 1620 cells, 833 actin foci. N = 3. **G**, Cells were double-transfected with Gal3-mOrange and anti-ORAI-1 siRNA, and subjected to vacuolar rupture imaging. SCRAMBLE: 30 SCVs, N = 3. siRNA ORAI-1: 45 SCVs. N = 4. Mann-Whitney. ns: not significant.

**Figure EV3.**
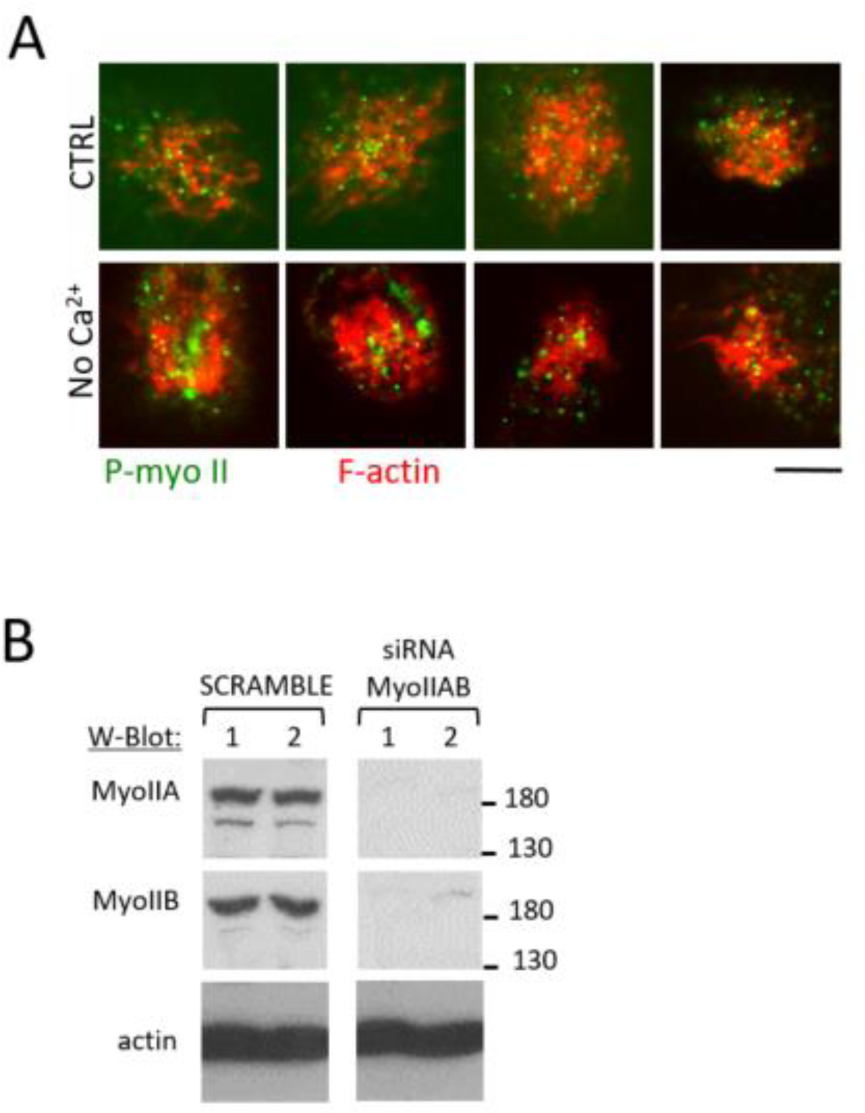
Role of extracellular Ca^2+^ on phospho-myosin II localization. **A**, Cells were challenged with *Shigella* for 15 min at 37°C in the presence (CTRL) or absence of extracellular Ca^2+^ (No Ca^2+^). Samples were fixed and processed for immunofluorescence staining of phospho-Myosin II (green) and F-actin (red). Representative fluorescence micrographs maximal of z-projections showing *Shigella*-induced actin foci. Scale bar = 5 μm. **B**, siRNA-mediated depletion of myosin IIA,B. Cells were transfected with the indicated siRNAs and processed for Western-blot analysis following SDS-PAGE in a 10 % polyacrylamide gel, using the antibody directed against the protein indicated on the left (W-Blot). Cells transfected for: 1: 24 hours; 2: 48 hours.

**Figure EV4.**
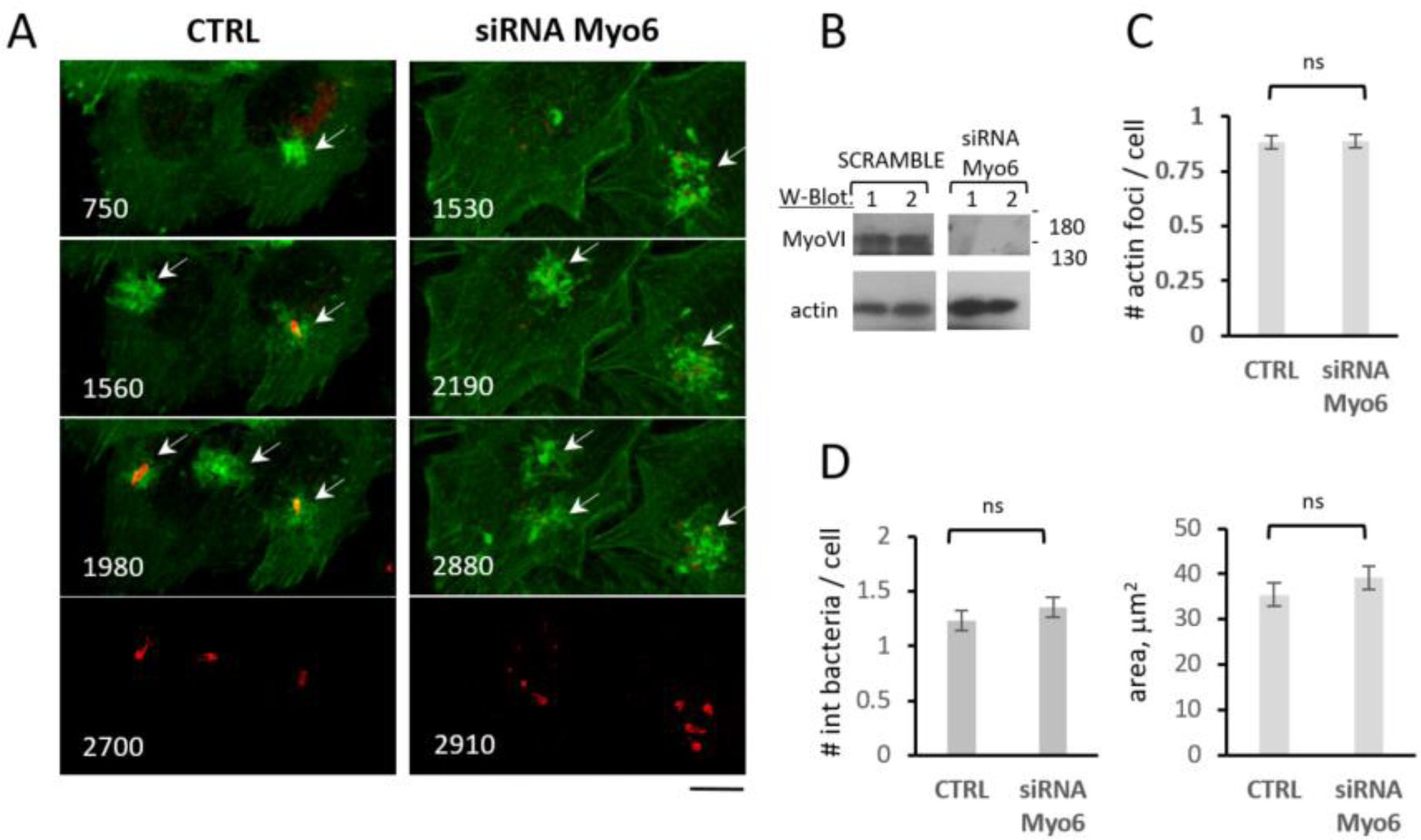
Myosin VI depletion does not affect *Shigella* invasion. Cells were triple - transfected with the indicated siRNA, Gal3-mOrange and GFP-actin, challenged with *Shigella*, and processed for actin foci and vacuolar rupture imaging. **A**, representative time-series of micrographs corresponding to maximal z-projections. Numbers: time relative to the start of actin foci formation in seconds. Red: Gal3-mOrange; Green: GFP-actin. The arrows point at actin foci. Scale bar = 5 μm. **B**, siRNA-mediated depletion of VI. Cells were transfected with the indicated siRNAs and processed for Western-blot analysis following SDS-PAGE in a 10 % polyacrylamide gel, using the antibody directed against the protein indicated on the left (W-Blot). Cells transfected with: 1: siRNA alone; 2: siRNA and GFP-actin. **C**, number of actin foci per cell. **D**, number of internalized bacteria based on SCV rupture events per cell. **E**, average area of actin foci. SCRAMBLE: 64 cells, 57 actin foci, N = 5. siRNA MyoVI: 57 cells, 51 actin foci, N = 5.

**Figure EV5.**
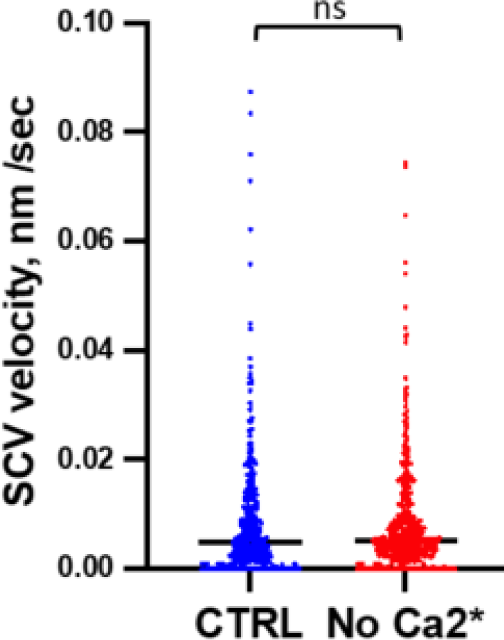
Effects of extracellular Ca^2+^ depletion on SCV motility. Cells were challenged with *Shigella* and SCVs were tracked for the first 300 seconds following rupture. SCV velocity in nm / sec. CTRL: control cells. No Ca2+: cells challenged in the absence of extracellular Ca^2+.^

## Expanded View Movies

**Movie EV1. Dual Ca^2+^ and vacuolar rupture imaging during *Shigella* invasion**. Green: Fluo-4 fluorescence. Red: Gal3-mOrange. Scale bar = 3 μm.

**Movie EV2. Dual GFP-actin and vacuolar rupture imaging of cells challenged with *Shigella* in control Ca^2+^ conditions.** Cells challenged in control conditions in the presence of 1.8 mM Ca^2+^. Red: Gal3-mOrange. Green: GFP-actin. Note the formation an actin coat around the SCV prior to rupture.

**Movie EV3. Dual GFP-actin and vacuolar rupture imaging of cells challenged with *Shigella* in the absence of extracellular Ca^2+^.** Cells challenged in the absence of extracellular Ca^2+^. Red: Gal3-mOrange. Green: GFP-actin. Note vacuolar rupture in the absence of detectable actin coat around the SCV prior to rupture.

**Movie EV4. Dual GFP-myosin VI and vacuolar rupture imaging of cells challenged with *Shigella*.** Red: Gal3-mOrange. Green: GFP-myosin VI. Note the formation of myosin VI patches interacting with the SCV.

**Movie EV5. Dual GFP-myosin VI and vacuolar rupture imaging of cells challenged with *Shigella* in the absence of extracellular Ca^2+^**. Red: Gal3-mOrange. Green: GFP-myosin VI. Note the lesser recruitment of myosin VI patches at the *Shigella* vacuole.

**Movie EV6. Dual GFP-actin and vacuolar rupture imaging of control cells challenged with *Shigella*. Left:** Red: Gal3-mOrange. Green: GFP-myosin VI. Right: Gal3-mOrange in gray levels. Time in seconds. Note the stability of Gal3-labeled *Shigella* vacuoles over several minutes, moving along the retrograde flow.

**Movie EV7. Dual GFP-actin and vacuolar rupture imaging of control cells challenged with *Shigella*. Left:** Red: Gal3-mOrange. Green: GFP-myosin VI. Right: Gal3-mOrange in gray levels. Time in seconds. Note the vacuolar membrane debris associated with actin comets.

**Movie EV8. Dual GFP-actin and vacuolar rupture imaging of anti-myosin VI siRNA treated cells challenged with *Shigella*. Left:** Red: Gal3-mOrange. Green: GFP-myosin VI. Right: Gal3-mOrange in gray levels. Time in seconds. Note the lesser stability and tumbling of Gal3-labeled *Shigella* vacuoles.

**Movie EV9. Dual GFP-actin and vacuolar rupture imaging of anti-myosin VI siRNA treated cells challenged with *Shigella*. Left:** Red: Gal3-mOrange. Green: GFP-myosin VI. Right: Gal3-mOrange in gray levels. Time in seconds. . Note the lesser stability and tumbling of Gal3-labeled *Shigella* vacuoles.

